# Integrative computational approach identifies new targets in CD4+ T cell-mediated immune disorders

**DOI:** 10.1101/2020.01.02.893164

**Authors:** Bhanwar Lal Puniya, Rada Amin, Bailee Lichter, Robert Moore, Alex Ciurej, Sydney Townsend, Ab Rauf Shah, Matteo Barberis, Tomáš Helikar

**Author notes:** These authors contributed equally to this work (Bailee Lichter, Robert Moore). **To whom correspondence should be addressed: Matteo Barberis**, Ph.D., Systems Biology, School of Biosciences and Medicine, Faculty of Health and Medical Sciences, University of Surrey, UK. or, **Tomáš Helikar**, Ph.D., Department of Biochemistry, University of Nebraska-Lincoln, USA.

## Abstract

CD4+ T cells provide adaptive immunity against pathogens and abnormal cells, and they are also associated with various immune related diseases. CD4+ T cells’ metabolism is dysregulated in these pathologies and represents an opportunity for drug discovery and development. Genome-scale metabolic modeling offers an opportunity to accelerate drug discovery by providing high-quality information about possible target space in the context of a modeled disease. Here, we develop genome-scale models of naïve, Th1, Th2 and Th17 CD4+ T cell subtypes to map metabolic perturbations in rheumatoid arthritis, multiple sclerosis, and primary biliary cholangitis. We subjected these models to *in silico* simulations for drug response analysis of existing FDA-approved drugs, and compounds. Integration of disease-specific differentially expressed genes with altered reactions in response to metabolic perturbations identified 68 drug targets for the three autoimmune diseases. *In vitro* experimental validations together with literature-based evidence showed that modulation of fifty percent of identified drug targets has been observed to lead to suppression of CD4+ T cells, further increasing their potential impact as therapeutic interventions. The used approach can be generalized in the context of other diseases, and novel metabolic models can be further used to dissect CD4+ T cell metabolism.

## Introduction

CD4+ T cells are essential components of the human immune system that fight against pathogenic invaders and abnormal cells by producing cytokines, and stimulating other cells, such as B cells, macrophages, and neutrophils.^1^ During immune response, CD4+ T cells are activated and proliferate, and their metabolism adjusts to fulfill increased bioenergetic and biosynthetic demands. For example, activated effector CD4+ T cells are highly glycolytic^2^ and use aerobic glycolysis and oxidative phosphorylation (OXPHOS) for proliferation.^3^ On the other hand, naïve, resting, and regulatory CD4+ T cells are less glycolytic and use OXPHOS and fatty acid oxidation (FAO) for energy generation. Accordingly, metabolically dysregulated CD4+ T cells were observed in several diseases such as diabetes,^4^ atherosclerosis,^5^ cancers,^6^ and autoimmune diseases such as rheumatoid arthritis (RA),^7,8^ multiple sclerosis (MS),^9^ primary biliary cholangitis (PBC),^10^ and systemic lupus erythematosus (SLE).^11,12^ Furthermore, metabolism of Type 1 T helper (Th1), Type 17 T helper (Th17), and inducible regulatory T cells have been found to be dysregulated in MS.^13^ Controlling CD4+ metabolic pathways can be important in fighting against some immune diseases. For example, CD4+ T cells are hyperactive in systemic lupus erythematosus (SLE), and inhibiting glycolysis as well as the mitochondrial metabolism improved the outcome in an animal model.^14^ Together, this evidence suggests a significant role of CD4+ T cell metabolism in immune-mediated diseases.

Repurposing existing drugs for novel indications represents a cost-effective approach for the development of new treatment options.^15^ Several studies have recently demonstrated the potential for drug repurposing in CD4+ T cell-mediated diseases.^16,17^ For example, 2-deoxy-D-glucose (anticancer agent) and metformin (antidiabetic drug) were shown to reverse SLE in a mouse model.^14^ However, drug repurposing, as well as drug discovery and development efforts for targeting T cell metabolism have been limited due the lack of knowledge about the key molecular targets in this context.

In recent years, analysis of large-scale biological datasets has emerged as a powerful strategy for discovery of novel mechanisms, drug targets, and biomarkers in human diseases.^18–21^ Here, we develop a computational modeling approach that integrates multi-omic data with systematic perturbation analyses of newly constructed whole-genome metabolic models of naïve CD4+ T cells, and Th1, Th2, and Th17 cells. This led to identification of potential drug targets for CD4+ T cell-mediated diseases (RA, MS, and PBC).

## Results

### Identification of genes expressed in the CD4+ T cells

We used the computational approach shown in Fig. 1 (see also *Supplementary Methods 1*) to construct metabolic models of naïve and effector CD4+ T cells. To identify metabolic genes expressed across CD4+ T cell subtypes (naïve, Th1, Th2, and Th17 cells), we integrated transcriptomics and proteomics data (*Supplementary Data 1*). The comparison of genes expressed in CD4+ T cell subtypes identified by different datasets are shown in *Supplementary Figure 1*. The analysis showed that between 675 and 836 metabolic genes were expressed depending on the CD4+ T cell subtype (*Supplementary Data 2*). Of these, 530 genes were expressed in all subtypes (Fig. 2a). On the other hand, 16, 25, 7, and 96 genes were specific to naïve, Th1, Th2, and Th17 cells, respectively. Pathway enrichment analysis using active metabolic genes suggested 6 enriched KEGG pathways common across all subtypes: carbon metabolism, TCA cycle, oxidative phosphorylation (OXPHOS), amino sugar and nucleotide sugar metabolism, and valine, leucine and isoleucine degradation (Fig. 2b). Fatty acid degradation and pentose phosphate pathway were enriched in naïve CD4+ T cells only, and fatty acid metabolism was enriched in the naïve, Th2, and Th17 subtypes. No specific KEGG pathways were found enriched solely in Th1, and Th17 cells. Among the enriched pathways shared by all CD4+ T cells, TCA cycle was enriched more than two-fold in naïve, Th1, and Th2 subtypes. Similarly, OXPHOS was enriched more than two-fold in naïve and Th1 subtypes (Fig. 2c). These results suggest that key metabolic pathways are active across all the subtypes. Importantly, the metabolism of various CD4+ T cell subtypes can be different with respect to these pathways’ levels of activity and the number of reactions active within the pathways.

**Fig. 1.**
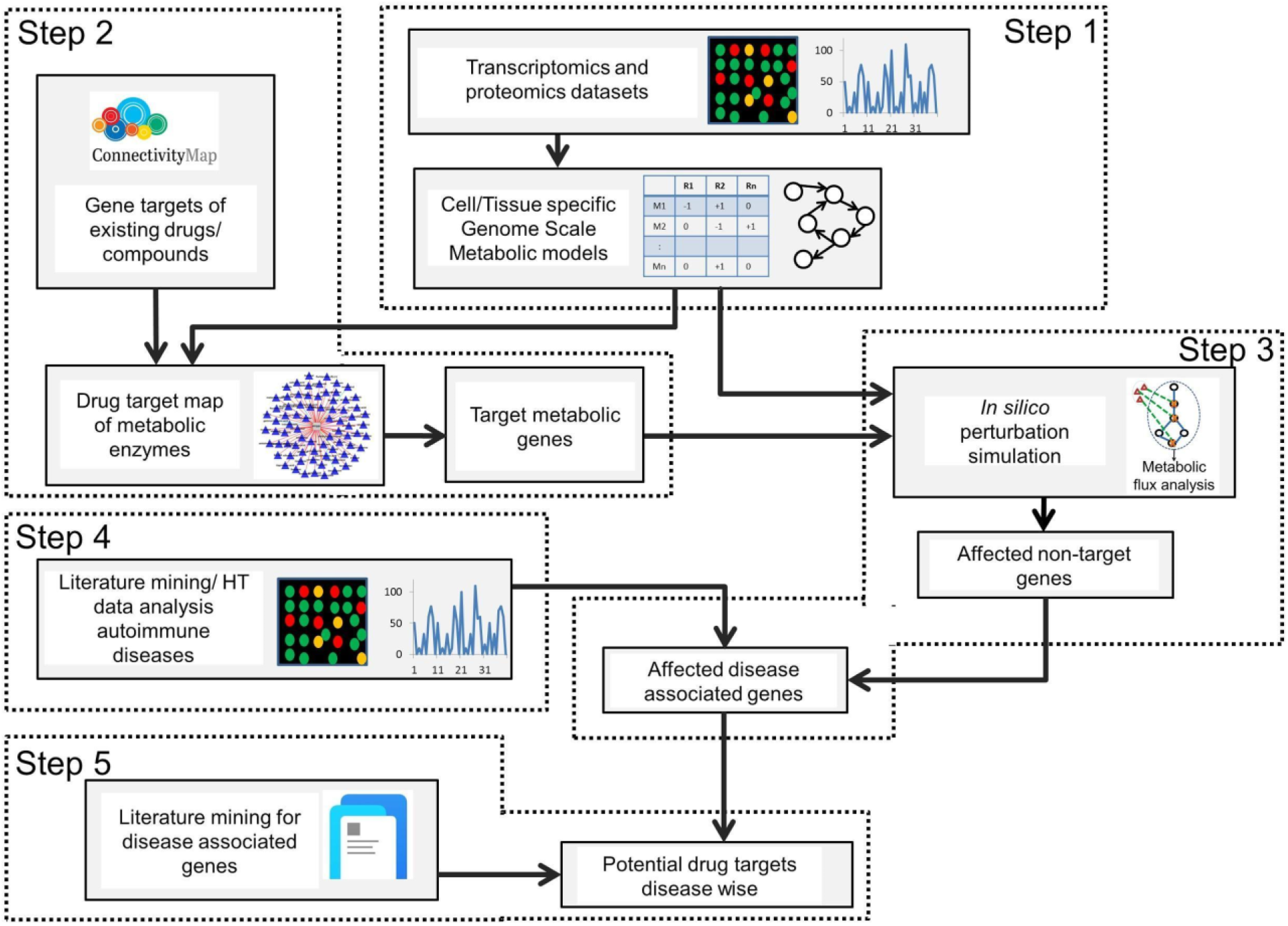
Integrative approach for the identification of potential metabolic drug targets. The computational approach comprised of five major steps: (1) Construction of metabolic models using integrated transcriptomics and proteomics data, (2) Identification of metabolic genes that are targets for existing drugs/compounds, (3) *In silico* inhibition of targets of existing drugs to identify affected reactions, (4) Identification of integrating differentially expressed genes (DEGs) in autoimmune diseases and integration with flux ratios obtained by perturbed and WT flux comparisons, and (5) Validation with literature and prediction of new targets.

**Fig 2.**
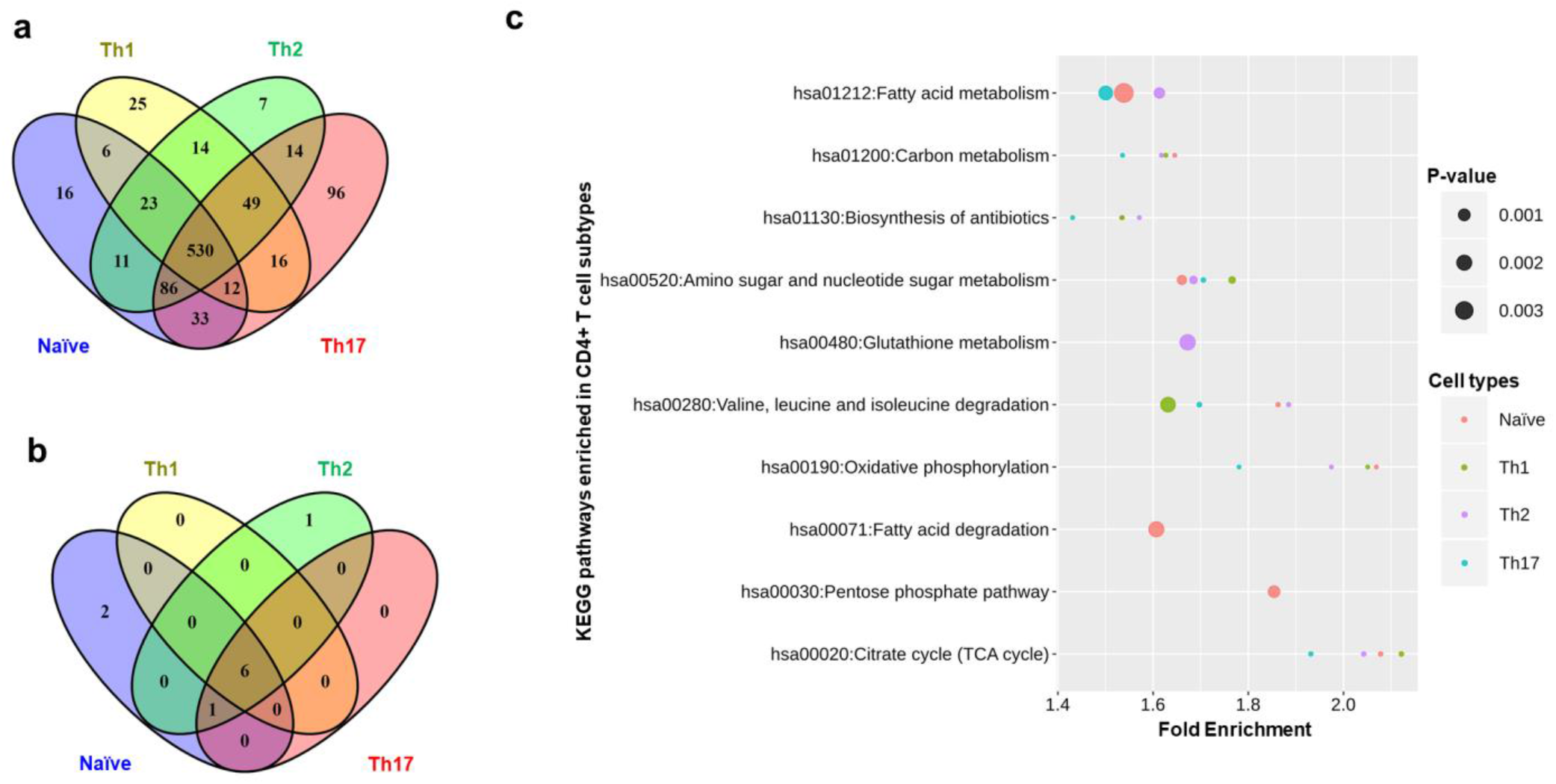
Construction of metabolic models in CD4+ T cells. (a) Active metabolic genes identified using integrated transcriptomics and proteomics data of CD4+ T cell subtypes. (b) KEGG pathway enrichment analysis of active genes in each cell type using all 1,892 metabolic genes as a background. (c) Fold enrichment and P-values (smaller sizes correspond to lower P-values) of KEGG pathways enriched across CD4+ T cell subtypes. A pathway was considered significantly enriched with P-value < 0.05 and false discovery rate (FDR) < 5%.

### Development and validation of genome-scale metabolic models of CD4+ T cells

To further examine these issues, we developed constraint-based metabolic models specific to naïve CD4+ T cells, Th1, Th2, and Th17 cells. Our genome-scale metabolic models comprised of 3956 to 5282 reactions associated with 1055 to 1250 genes (Table 1; *Supplementary Dataset 1*). The number of internal enzyme-catalyzed reactions were 2501, 1969, 2549, and 2640 for naïve, Th1, Th2, and Th17 models respectively, distributed across 84 metabolic pathways (*Supplementary Figure 2; note that transport and exchange reactions were excluded*). The models include more genes associations than active genes identified from the data because the model-building algorithm inserts some reactions that are not supported by data but required for the model to achieve essential metabolic functions for biomass production (see *Materials and Methods*). We compared our models with the existing model for naïve CD4+ T cell (CD4T1670) (Table 1). Since our models exclude blocked reactions and dead ends, we have removed dead ends from CD4T1670 for fair comparisons. Since we based all models of this study on the more recent human metabolic network Recon3D, they include more reactions and metabolites than CD4T1670, based on Recon2.

**Table 1:** Metabolic models of CD4+ T cells

|  | Naïve | Th1 | Th2 | Th17 | CD4T1670<br>(Naïve)<br>(reduced)* |
| --- | --- | --- | --- | --- | --- |
| **Genes** | 1055 | 1133 | 1127 | 1250 | 1027 |
| **Reactions** | 5179 | 3956 | 5252 | 5282 | 2592 |
| **Internal reactions**<br>**(Enzyme catalyzed)** | 2501 | 1969 | 2549 | 2640 | 1146 |
| **Metabolites** | 3153 | 2517 | 3156 | 3263 | 1402 |

\* Our models do not include dead ends and use gene IDs. To make CD4T1670 comparable to our models we removed dead ends and counted the number of genes in the reduced model (in place of transcripts).

We validated the models based on the active pathways and gene essentiality. We first identified pathways that are known to be active in different CD4+ T cell subtypes (see *Materials and Methods*) and searched for their activity (with non-zero fluxes) in the corresponding models through Flux Balance Analysis (FBA). Several major pathways were in agreement with the literature. These include glycolysis, TCA cycle, glutaminolysis and pyruvate to lactate conversion (aerobic glycolysis) that showed non-zero flux in all the models. We present an illustrative flux map of the pathways mentioned above for the naïve model in Fig. 3. The figure also indicates flux differences with the Th1 model for critical reactions. Flux maps for the Th1, Th2, and Th17 models are shown in *Supplementary Figure 3, 4*, and *5*. Furthermore, we show some specific observed behaviors of CD4+ T cells collected from the literature used for validation in Table 2. For comparison, we also performed these validations in CD4T1670, also shown in Table 2. In all the models developed in this study, limiting glucose from the environment resulted in a decreased growth rate (Fig. 4a), in agreement with existing knowledge but not reproduced by CD4T1670 model (*Supplementary Figure 6*). All the models produced lactate (Table 2, *Supplementary Figure 7*). Literature shows that increasing PDHm (pyruvate dehydrogenase) by inhibiting PDHK (pyruvate dehydrogenase kinase) would redirect flux from pyruvate to TCA cycle and, therefore, will decrease the lactate production. This was reproduced by our models (Figure 4b). Our models were also able to reproduce the essentiality of Leucine, Arginine, and ACC1 for T cell growth (Table 2), in agreement with the literature. It’s important to note that the activity of some pathways in the models was not in agreement or partially agreeing with the literature. Specifically, we did not observe a significant effect on growth rate when glutamine^22^ was removed from exchange reactions in the effector CD4+ T cell models (Fig. 4c). However, literature has shown that while transporters of glutamine in activated CD4+ T cells are dispensable, removal of glutamine is critical and affects growth. Our systematic analyses showed that (1) inhibiting glutamine synthase (GLNS) (that converts glutamate to glutamine) in the absence of glutamine uptake in the model decreases growth to zero, and (2) glutamine can have an impact on growth when glucose availability is limited (less than 2 mmol/g.DW/hr) (Fig. 4d). Using CD4T1670, no effect on biomass was observed when glutamine was varied in presence and absence of glucose (*Supplementary Figure 8*). Thus, we may hypothesize that glutamine might be conditionally critical for CD4+ T cells specifically when availability of other nutrients is low.

**Fig. 3.**
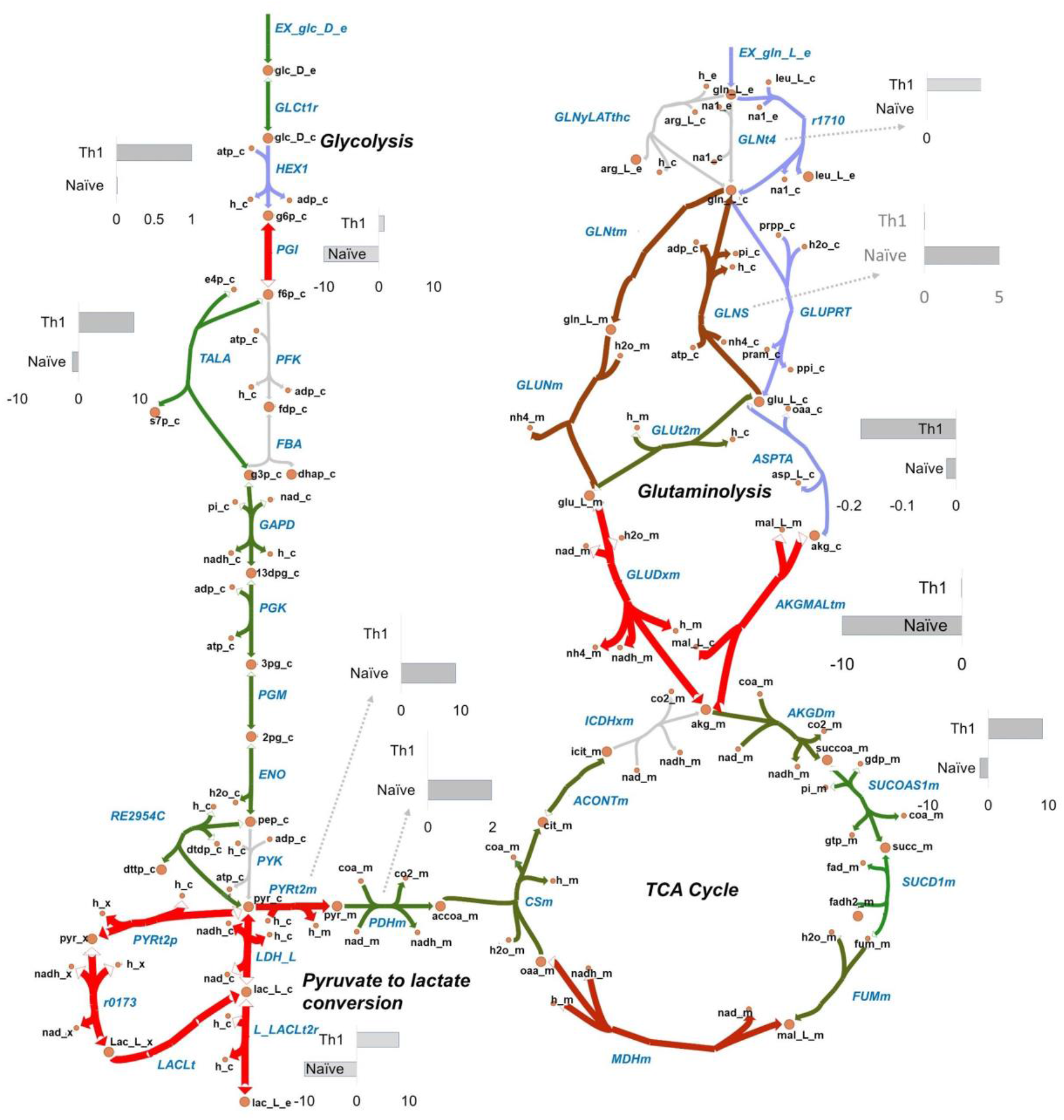
Flux maps of metabolic pathways active in CD4+ T cell metabolic models. Escher maps showing fluxes through glycolysis, glucose to lactate conversion, TCA cycle, glutaminolysis in naïve (a) and Th1 (b) models. Both naïve and Th1 models convert pyruvate to lactate (aerobic glycolysis). In glycolysis, the naïve model had the reverse direction flux through PGI reaction while Th1 cells have forward direction flux. All the models exhibit an uptake of glutamine that ultimately forms α-Ketoglutaric acid (glutaminolysis). GLNtm (glutamine transporter) and GLUNm (convert glutamine to glutamate) reactions are active in naïve model and not in Th1 model that use different routes for glutamine to glutamate conversion.

**Fig. 4:**
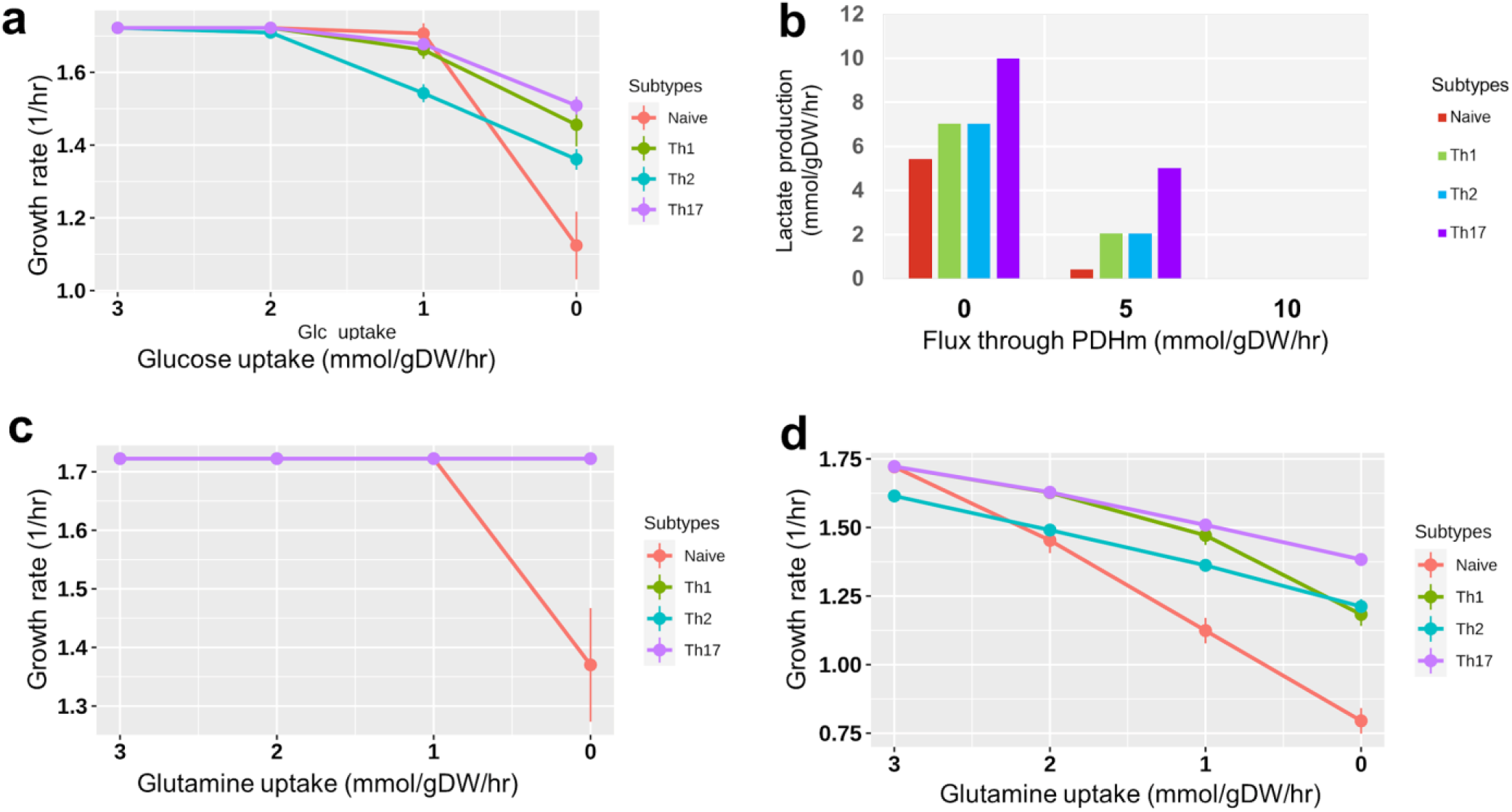
Validation of metabolic models. (a) Dependency of growth rate to varying rate of glucose uptake (b) Production of lactate with increased flux through pyruvate dehydrogenase. (c) The dependency of growth rate (in all models) on glutamine when glucose was available (> 5 mmol/ gDW/hr). (d) The dependency of growth on glutamine when glucose was removed from the environment. Dots in Figure (a, c, and d) are average flux and error bars are standard deviation (n = 5).

**Table 2:** Model validation using specific behaviors

| Observation from literature | *In silico* experiment | Naïve | Th1 | Th2 | Th17 | CD4T1670 (Naïve) |
| --- | --- | --- | --- | --- | --- | --- |
| 1. Glucose transporter deficiency will inhibit the proliferation of cells.<sup>74,91</sup> | Inhibited glucose transporters / remove glucose from the environment | ↓growth | ↓growth | ↓growth | ↓growth | No change |
| 2. CD4+ T cells produce lactate.<sup>45</sup> | Observed if models have produced lactate | Lactate + | Lactate + | Lactate + | Lactate + | Lactate + |
| 3. Activated CD4+ T cells control pyruvate dehydrogenase function which in turn redirect pyruvate from entering the TCA cycle towards lactate production.<sup>91</sup> | Increasing flux through pyruvate dehydrogenase reaction (PDHm) should decrease the lactate production | ↓Lactate | ↓Lactate | ↓Lactate | ↓Lactate | NA |
| 4. During T cell activation GAPDH is important for glycolysis.<sup>91</sup> | Investigating if inhibition of GAPDH is impacting the growth | ↓growth | 0 growth | 0 growth | 0 growth | ↓growth |
| 5. Lack of transporters for Leucine impairs the metabolic reprogramming required Th1 and Th17.<sup>91</sup> | Remove leucine from media and investigate the impact on growth | 0 growth | 0 growth | 0 growth | 0 growth | 0growth |
| 6. In T cells arginine deprivation leads to cell cycle arrest.<sup>91</sup> | Remove arginine from the media and investigate the impact on growth | 0 growth | 0 growth | 0 growth | 0 growth | 0growth |
| 7. Gene acetyl-CoA carboxylase alpha (ACC1) is essential for T cell homeostasis.<sup>91</sup> | Inhibit gene ACC1 or reaction Acetyl Coenzyme A Carboxylase (ACCOAC) and investigate the impact on growth | No change | 0 growth | 0 growth | 0 growth | No change |
| 8. Glutamine deficiency in media leads to low growth and proliferation.<sup>91</sup> | Removing glutamine from media | ↓growth | No change | No change | No change | No change |

\* ↓ = decrease, + = produced, light green = agreed with literature, gray = partial agreement, red = disagreement, NA = experiment was not executed because infeasible.

Next, we predicted essential genes, and compared the results against independent data to identify overlap between genes predicted by our models and identified from experiments. Because large scale essentiality datasets were not available for CD4+ T cells, we used essentiality datasets available for different human cancer cell lines. Gene deletion analysis predicted 84, 95, 81, and 84 genes as essential in the naïve, Th1, Th2, and Th17 models respectively (*Supplementary Data 3*). More than 70% of these predictions agreed with genes experimentally defined as essential and conditionally essential in those different but related cell lines.^23^ (*Supplementary Figure 9*). To assess if higher ranked predictions compared better with gene essentiality data, we generated precision-recall curves using 84, 95, 81, and 84 genes respectively from each model that were identified as essential. The area under curve for all models was > 75% (*Supplementary Figure 10 - 13*). Additional validations based on CD4+ T cell-specific essential functions are presented in *Supplementary Methods 3*. Overall, the validation confirmed that our constraint-based metabolic models specific to naïve CD4+ T cells, Th1, Th2, and Th17 cells represent relevant and realistic systems to examine drug response and predict drug targets.

### Mapping existing, and identifying potential drug targets in CD4+ T cells

We used the validated CD4+ T cell-specific models to predict potential drug targets and combined it with the publicly available drug repurposing and tool compound data set from the Connectivity Map (cMap) database and mapped the approved drugs, clinical drug candidates and tool compounds in the dataset with the metabolic genes in the models (Fig. 5a). Next, we performed *in silico* knock-outs of the associated drug target genes. Due to the presence of isozymes, not all the deleted target genes influenced the reaction(s). We identified 86, 79, 86, and 90 target genes whose deletion blocks at least one associated reaction in naïve, Th1, Th2, and Th17 models, respectively (Fig. 5b). In turn, these disruptions block multiple downstream reactions. Of these, 62 were common among four CD4+ T cell subtypes (Fig. 5c). Four genes were targeted only in Th1 cells. All modeled gene deletions resulted in altered flux distributions that were quantified using flux ratios. For each drug target deletion, we classified all reactions into three categories (see *Materials and Methods*): (1) reactions with decreased fluxes (down-reactions), (2) reactions with increased fluxes (up-reactions), and (3) reactions without any changes. We used these flux ratios to identify potential drug targets specific for immune diseases, by exploring how disease-specific metabolic functions are affected upon each drug target inhibition.

**Fig. 5:**
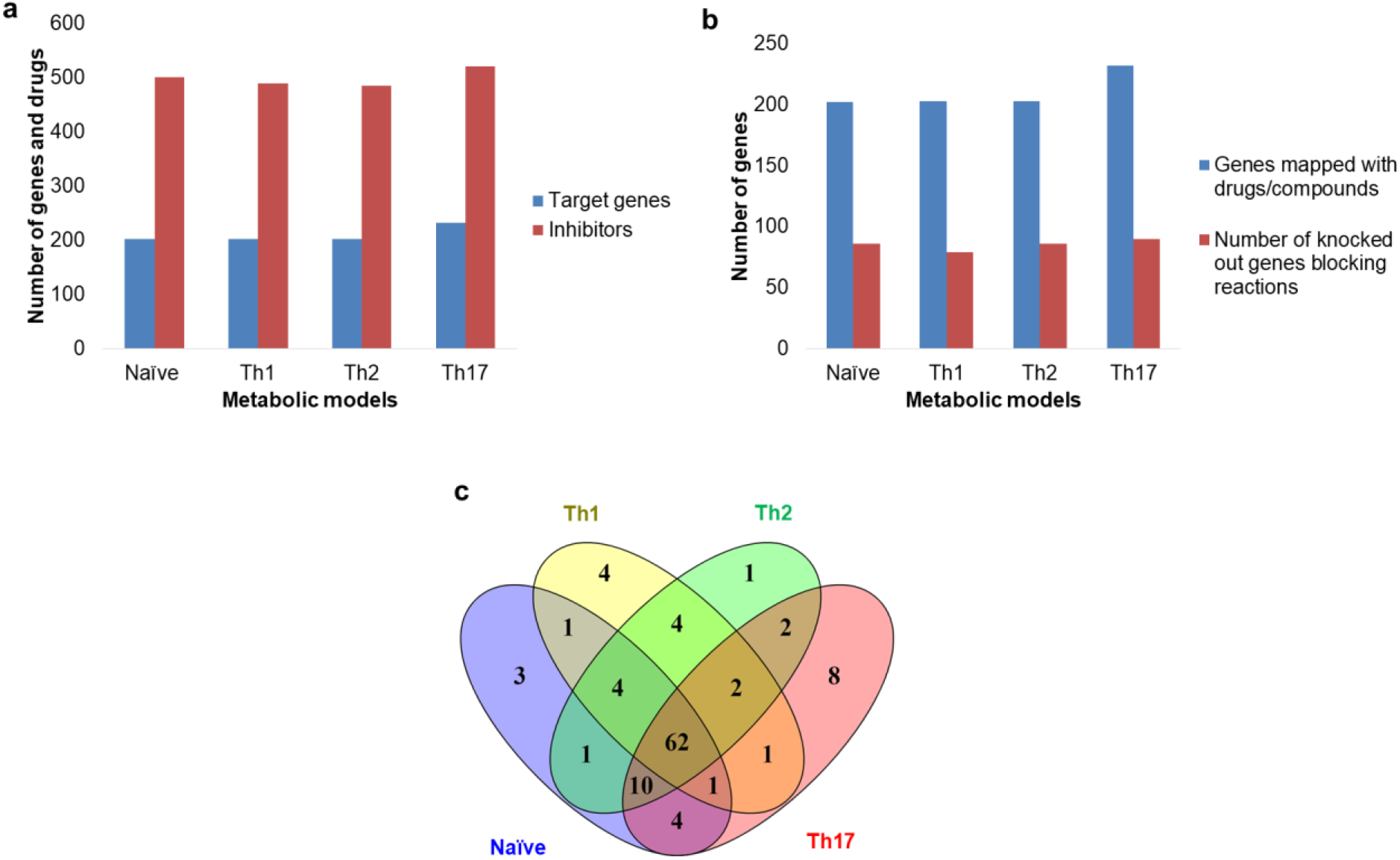
Drug targets in CD4+ T cell models. (a) Distribution of metabolic drug target genes, and inhibitory drugs or compounds in each model. (b) Number of metabolic genes in the models mapped with inhibitory drugs (blue bars) and number of genes among drugs mapped genes that can block at least one reaction upon inhibition (red bars). (c) Comparison of metabolic drug targets that affect reactions upon deletion in CD4+ T cell models.

First, we identified disease-specific metabolic functions for RA, MS, and PBC using differential gene expression analysis of publicly available patients’ data (Case-Control studies) (see *Materials and Methods*). We identified 852, 1459, and 553 differentially expressed genes (DEGs) for RA, MS, and PBC, respectively (*Supplementary Data 4*). From these DEGs, we selected genes relevant to our metabolic models. For example, 36 metabolic genes were upregulated and 27 genes were downregulated in RA (Fig. 6a). Biological process enrichment analysis identified purine metabolism, and starch sucrose metabolism as enriched in upregulated genes. On the contrary, lysine degradation, fatty acid elongation, and carbon metabolism were downregulated. Enriched metabolic pathways for all three diseases are shown in *Supplementary Data 4*.

**Fig. 6:**
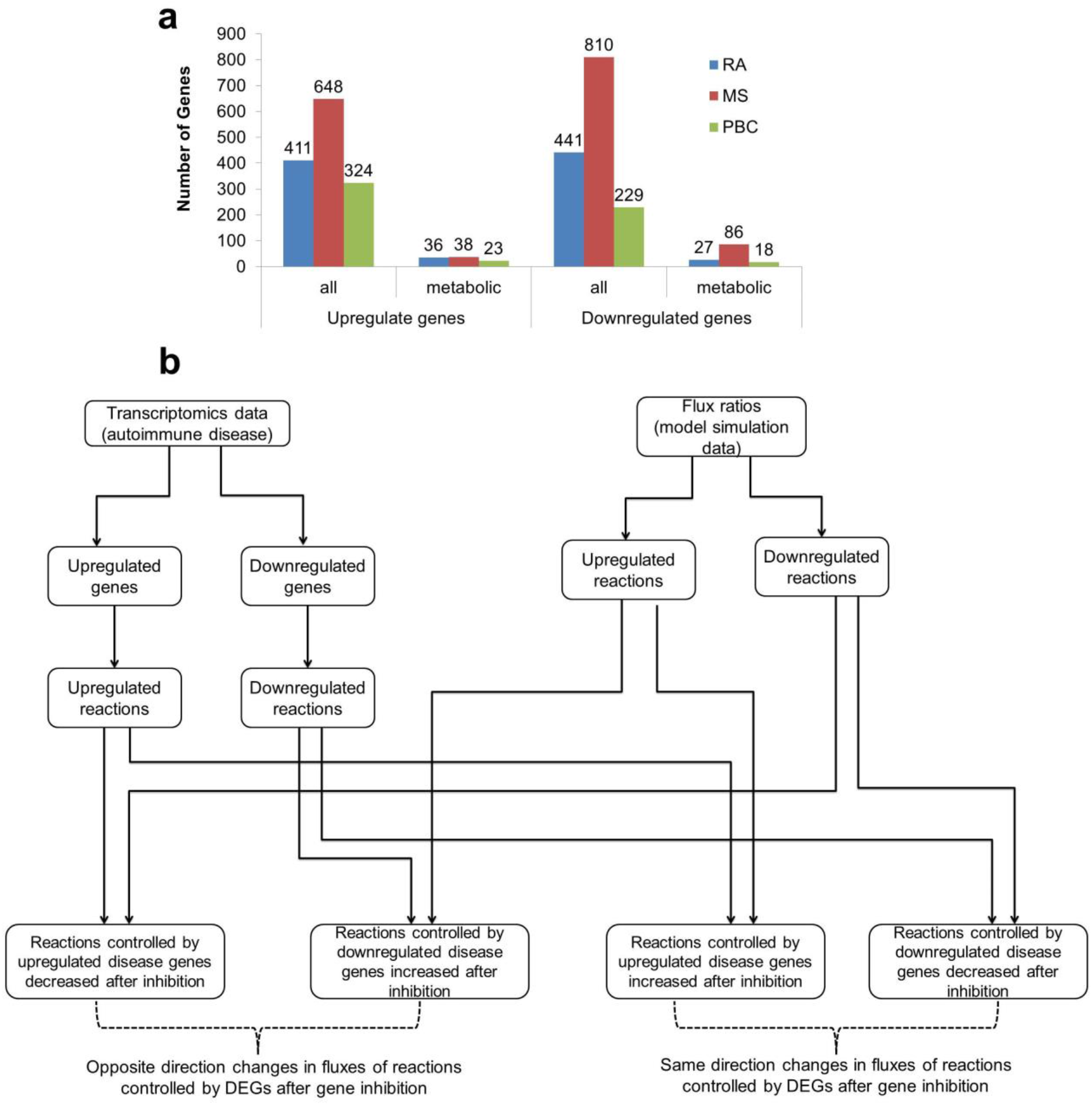
Identification of potential drug targets for RA, MS, and PBC. (a) Number of all differentially expressed genes (DEGs) and metabolic DEGs in three diseases rheumatoid arthritis (RA), multiple sclerosis (MS), and primary biliary cholangitis (PBC). The DEGs were analyzed using three transcriptomics datasets (one dataset per disease). The data were obtained from peripheral CD4+ T cells of groups of patients and healthy individuals. (b) Schematic representation of the integration of disease-associated differentially expressed genes and affected reaction on each drug target gene perturbation. For each drug target deletion, we investigated how many of fluxes regulated by upregulated genes are decreased and fluxes regulated by downregulated by increased. We used these numbers to calculate PES (perturbation effect score, *see Materials and Methods*).

To identify potential drug targets for the aforementioned diseases, we looked for target genes whose deletion (inhibition) would have the appropriate effect on diseases’ DEGs. For each gene inhibition, we specifically investigated the decrease in metabolic flux through reactions controlled by genes upregulated in disease, and increase in metabolic flux through reactions controlled by genes downregulated in disease (Fig. 6b). Using flux ratios of metabolic DEGs, we calculated a *perturbation effect score* (PES; see *Materials and Methods*) for each drug target gene in each pair model/disease. PES represent the effect of gene inhibition on both upregulated and downregulated genes. A positive PES value for the drug target gene means that its inhibition decreases more fluxes controlled by genes upregulated in disease than it increases or increases more fluxes controlled by genes downregulated in disease than it decreases. As such, inhibition of that gene target reverses the fluxes controlled by disease DEGs. In contrast, a negative PES means that the inhibition of a target gene increases more fluxes controlled by upregulated genes or decrease the more fluxes controlled by downregulated genes than the opposite. Among the different combinations of cell types and diseases, the PESs range was from −2 to 2 (*Supplementary Figure 14*). Based on these considerations, genes with higher positive PES can serve as potential drug targets for the disease.

Using PES as a measure of target relevance, we identified 62 potential drug targets that were common to our models (Fig. 5c). These genes displayed various PES ranks across models and diseases. To choose drug targets that performed better across different CD4+ T cells, we considered PES ranks of the four subtype-specific models. First, we normalized the PES ranks by transforming them to Z-scores in each model. Since the studied diseases typically involve more than one type of CD4+ T cell subtype, we next summed up the Z-scores of all the models within a disease for each drug target (*Supplementary Figure 14*). A minimum aggregate Z-score represents overall high PES ranks predicted across four cell types. Therefore, a gene with a minimum aggregated Z-score could be a potential high confidence drug target. We used a Z-score cutoff of −1 (1 standard deviation lower than the mean aggregated Z-score) and identified 17, 27, and 24 potential drug targets for RA, MS, and PBC, respectively (Table 3). Ranking based on aggregated Z-scores is provided in *Supplementary Data 5*. Taken together, our combined use of the disease-matched genome-scale metabolic models of CD4+ T cells and the well target-annotated public dataset of bioactive compounds generated a manageable list of potential drug targets suitable for deeper analysis and follow-up.

**Table 3:**
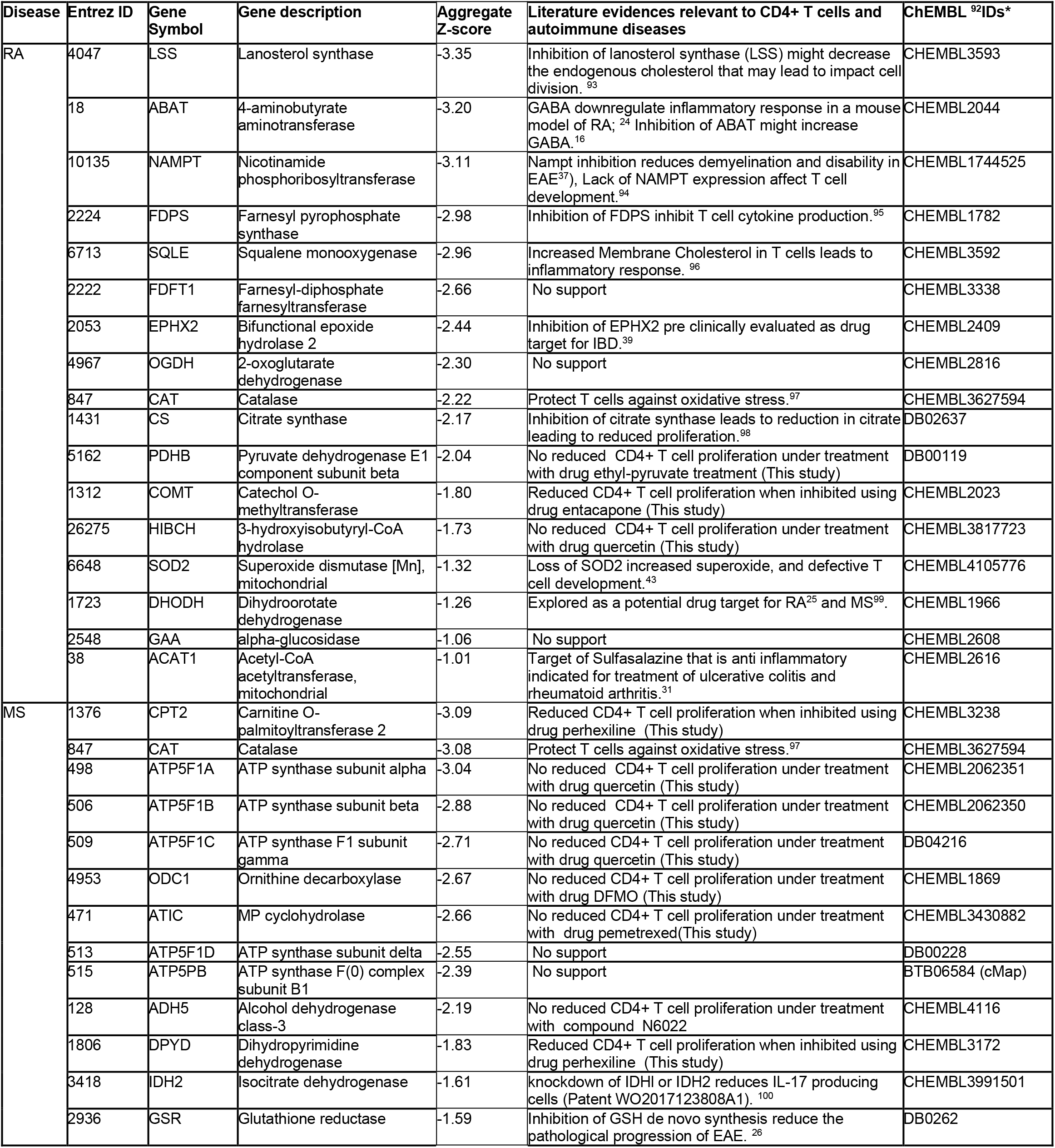
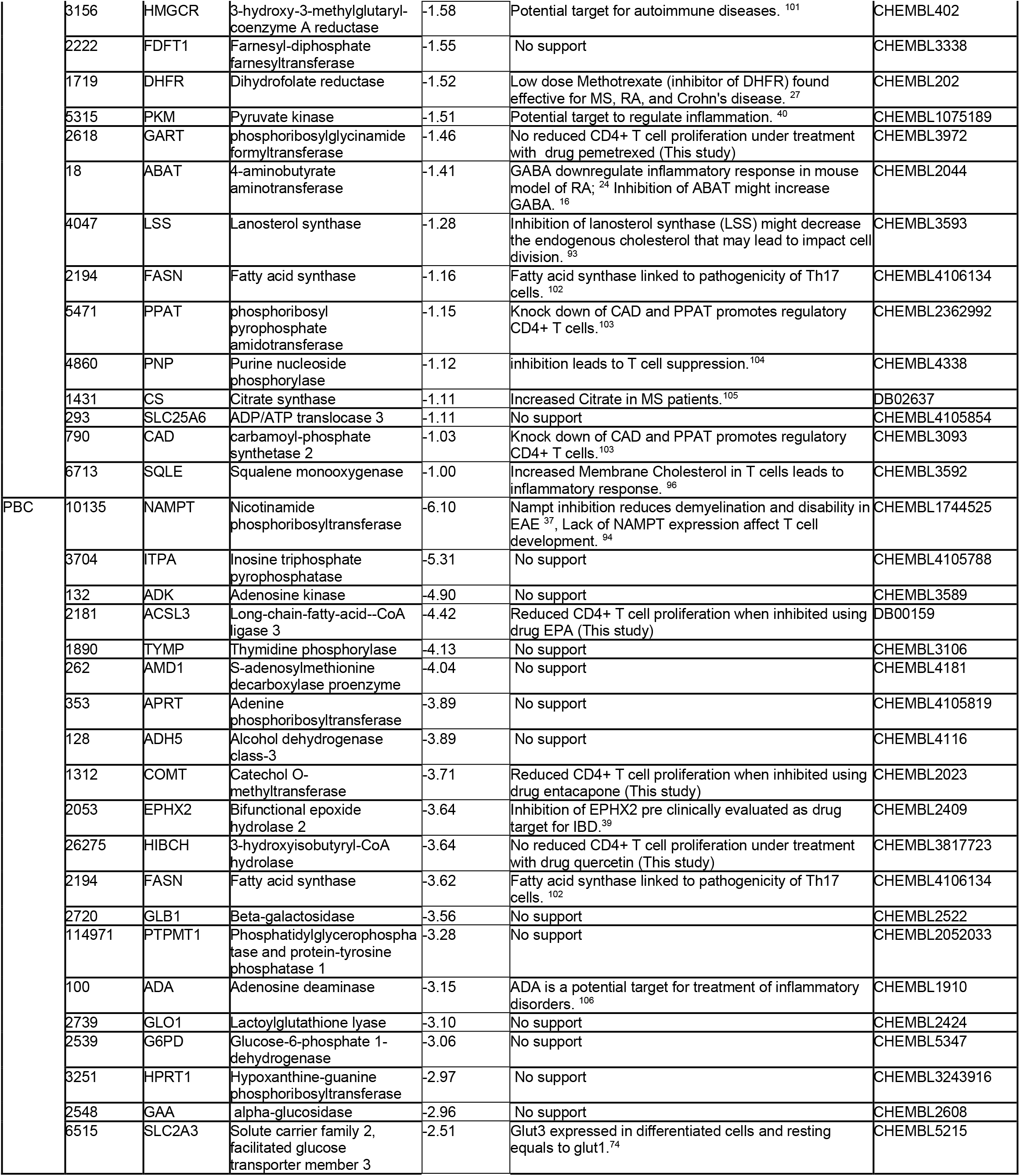
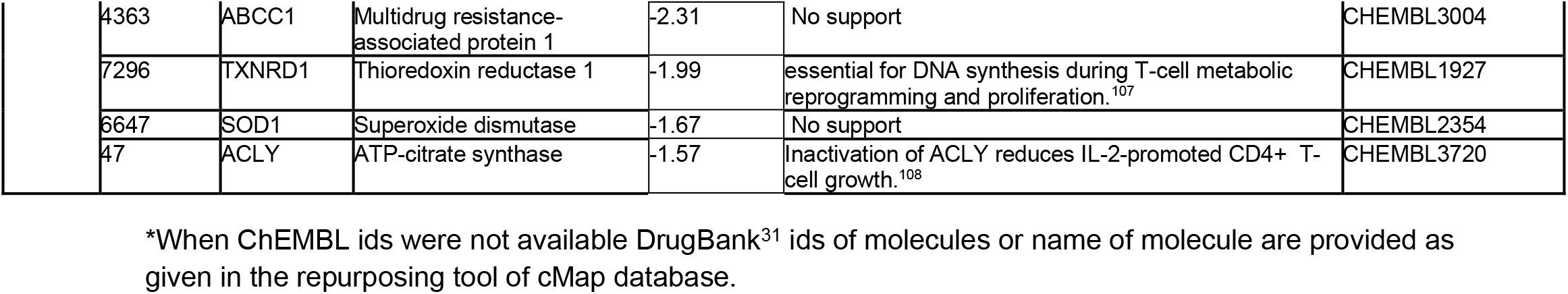
Identified CD4+ T cell drug targets for RA, MS, and PBC

### Analysis and validation of predicted drug targets

To further analyze and validate our target list, we performed a comprehensive literature survey (Table 3). Among the 17 suggested drug targets for RA, dihydroorotate dehydrogenase (DHODH) and Acetyl-CoA acetyltransferase (ACAT1) have been already explored as targets in drug development efforts^24,25^, and 15 genes were newly identified. Among these, eight (LSS, NAMPT, FDPS, SQLE, EPHX2, CAT, CS, SOD2) have been found to inhibit CD4+ T cell proliferation upon deletion (Table 3). The product of the reaction catalyzed by 4-Aminobutyrate Aminotransferase (ABAT) is linked to RA. Dysregulation of other genes, such as pyruvate dehydrogenase E1 (PDHB), Farnesyl-diphosphate farnesyltransferase 1 (FDFT1), Oxoglutarate Dehydrogenase (OGDH), alpha-galactosidase (GAA), has not been previously reported to impact CD4+ T cell proliferation.

Furthermore, we predicted 27 possible drug targets for MS. Of these, glutathione reductase (GSR), and dihydrofolate reductase (DHFR) were already explored as targets using the experimental autoimmune encephalomyelitis (EAE) model^26,27^ and 25 genes were newly identified. Among these, 12 (CAT, IDH2, HMGCR, PKM, ABAT, LSS, FASN, PPAT, PNP, CS, CAD, SQLE) have been previously reported inhibiting CD4+ T cell proliferation upon deletion.

Genes that were not previously reported to affect CD4+ T cells upon deletion include Carnitine O-palmitoyltransferase 2 (CPT2), MP cyclohydrolase (ATIC), Ornithine decarboxylase (ODC1), Dihydropyrimidine dehydrogenase (DPYD), and Farnesyl-diphosphate farnesyltransferase (FDFT1).

Finally, we identified 24 possible drug targets for PBC. None of them was previously explored as a drug target in PBC. Deletion of seven of these potential gene targets (NAMPT, EPHX2, FASN, ADA, SLC2A3, TXNRD1, ACLY) has been reported to affect CD4+ T cells in the literature. Genes that have not yet been reported to affect CD4+ T cells upon deletion include Long-chain-fatty-acid--CoA ligase 3 (ACSL3), Adenosine kinase (ADK), and S-adenosylmethionine decarboxylase proenzyme (AMD1).

43 of the 68 predicted drug targets were found for only one disease. Six drug targets (LSS, ABAT, SQLE, FDFT1, CAT, CS) were common to RA and MS; five drug targets (NAMPT, EPHX2, COMT, HIBCH, GAA) were in common between RA and PBC; and two drug targets (ADH5, FASN) were in common between MS and PBC. We show the drugs and compounds available for these targets in Table 4. Out of the 55 unique drug targets identified across three diseases, 38 were found to be robust across different cutoffs (Supplementary method 9). A few examples of drug targets from purine metabolism, fatty acid synthesis, and TCA cycle are shown in Fig. 7.

**Fig. 7:**
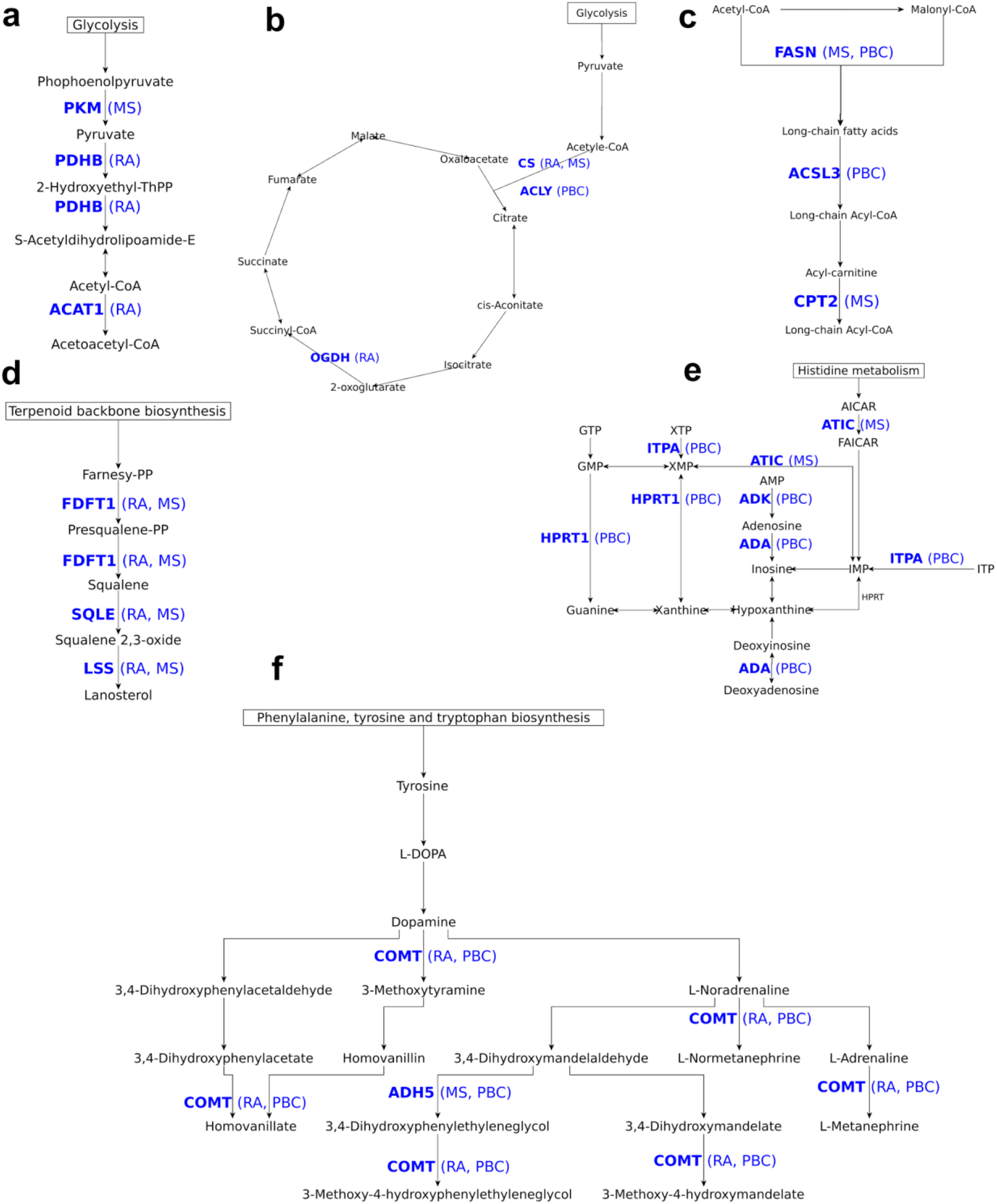
Examples of identified drug targets mapped on the metabolic pathways. Relevant sub-networks of pathways where drug targets were mapped are shown for (a) pyruvate metabolism, (b) TCA cycle, (c) fatty acid biosynthesis, (d) steroid biosynthesis, (e) purine metabolism, and (f) tyrosine metabolism. The mapped drug targets (bold font) and diseases (in brackets) are shown in blue colored text.

**Table 4:** Drugs and compounds for identified drug targets

| Gene Symbol | Gene description | Drugs/ compound* | Status of drugs/Compounds* |
| --- | --- | --- | --- |
| LSS | Lanosterol synthase | R0-48-8071 | Preclinical |
| ABAT | 4-aminobutyrate aminotransferase | Vigabatrin, Phenelzine, valproic acid | launched |
| NAMPT | Nicotinamide phosphoribosyltransferase | FK866 | Phase 2 |
| FDPS | Farnesyl pyrophosphate synthase | Pamidronic acid, Zoledronic acid, Alendronic acid, Ibandronate, Risedronic acid | Launched |
| SQLE | Squalene monooxygenase | Ellagic-acid | Phase 2 |
| FDFT1 | Farnesyl-diphosphate farnesyltransferase | TAK-475 | Investigational |
| EPHX2 | Bifunctional epoxide hydrolase 2 | GSK2256294A | Phase 1 |
| OGDH | 2-oxoglutarate dehydrogenase | Valproic acid | Launched |
| CAT | Catalase | Fomepizole | Launched |
| CS | Citrate synthase | Oxaloacetate | Phase 2/ Phase 3 |
| PDHB | Pyruvate dehydrogenase E1 component subunit beta | 2-oxopropanoate | Preclinical |
| COMT | Catechol O-methyltransferase | Entacapone, Nitecapone, Opicapone | Launched, Phase 2, phase 3 |
| HIBCH | 3-hydroxyisobutyryl-CoA hydrolase | Quercetin | Launched |
| SOD2 | Superoxide dismutase [Mn], mitochondrial | Tetraethylenepentamine | Phase 2/ Phase 3 |
| DHODH | Dihydroorotate dehydrogenase | Atovaquone, Leflunomide, Teriflunomide, Brequinar | Launched |
| GAA | alpha-glucosidase | Miglitol, Acarbose | Launched |
| ACAT1 | Acetyl-CoA acetyltransferase, mitochondrial | Sulfasalazine | Launched |
| CPT2 | Carnitine O-palmitoyltransferase 2 | Perhexiline | Launched |
| ATP5F1A | ATP synthase subunit alpha | Quercetin | Launched, |
| ATP5F1B | ATP synthase subunit beta | Quercetin | Launched |
| ATP5F1C | ATP synthase F1 subunit gamma | Quercetin | Launched |
| ODC1 | Ornithine decarboxylase | MC-1, Putrescine, DFMO | Phase 3, Phase 2, launched |
| ATIC | MP cyclohydrolase | Pemetrexed | Launched |
| ATP5F1D | ATP synthase subunit delta | Sevoflurane, Enflurane, Methoxyflurane | Launched |
| ATP5PB | ATP synthase F(0) complex subunit B1 | BTB06584 | Preclinical |
| ADH5 | Alcohol dehydrogenase class-3 | N6022 | Phase 1/Phase 2 |
| DPYD | Dihydropyrimidine dehydrogenase | 5-fluorouracil, Gimeraci | Launched |
| IDH2 | Isocitrate dehydrogenase | AGI-6780 | Preclinical |
| GSR | Glutathione reductase | Carmustine | Launched |
| HMGCR | 3-hydroxy-3-methylglutaryl-coenzyme A reductase | Atorvastatin, Fluvastatin, Lovastatin, Meglutol, Pitavastatin, Pravastatin, Rosuvastatin, Simvastatin, Nadide | Launched |
| DHFR | Dihydrofolate reductase | Aminopterin, Chlorproguanil, Methotrexate, Pralatrexate, Proguanil, Pyrimethamine, Sulfameter, Trimethoprim, Pemetrexed | Launched |
| PKM | Pyruvate kinase | TEPP-46, 2-oxopropanoate | Preclinical |
| GART | phosphoribosylglycinamide formyltransferase | Pemetrexed | Launched |
| FASN | Fatty acid synthase | Pyrazinamide, Cerulenin | Launched |
| PPAT | phosphoribosyl pyrophosphate amidotransferase | Azathioprine, Mercaptopurine | Launched |
| PNP | Purine nucleoside phosphorylase | Acyclovir, Didanosine | Launched |
| SLC25A6 | ADP/ATP translocase 3 | Clodronic-acid | Launched |
| CAD | carbamoyl-phosphate synthetase 2 | Sparfosate | Phase 3 |
| ITPA | Inosine triphosphate pyrophosphatase | Citric-acid | Preclinical |
| ADK | Adenosine kinase | ABT-702 | Preclinical |
| ACSL3 | Long-chain-fatty-acid--CoA ligase 3 | Eicosapentaenoic Acid | Launched |
| TYMP | Thymidine phosphorylase | Tipiracil | Launched |
| AMD1 | S-adenosylmethionine decarboxylase proenzyme | Ademetionine, Putrescine | Launched, Phase 2 |
| APRT | Adenine phosphoribosyltransferase | Citric-acid | Preclinical |
| GLB1 | Beta-galactosidase | Fagomine | Phase 2 |
| PTPMT1 | Phosphatidylglycerophosphatase and protein-tyrosine phosphatase 1 | Alexidine | Preclinical |
| ADA | Adenosine deaminase | Cladribine, Pentostatin, Dipyridamole, Fludarabine | Launched |
| GLO1 | Lactoylglutathione lyase | Indomethacin | Launched |
| G6PD | Glucose-6-phosphate 1-dehydrogenase | RRx-001 | Phase 2 |
| HPRT1 | Hypoxanthine-guanine phosphoribosyltransferase | Azathioprine, Mercaptopurine | Launched |
| SLC2A3 | Solute carrier family 2, facilitated glucose transporter member 3 | 2-deoxyglucose | Phase 2 |
| ABCC1 | Multidrug resistance-associated protein 1 | Reversan, Ko143 | Preclinical |
| TXNRD1 | Thioredoxin reductase 1 | Fotemustine | Launched |
| SOD1 | Superoxide dismutase | Tetraethylenepentamine | Phase 2/Phase 3 |
| ACLY | ATP-citrate synthase | ETC-1002 | Phase 3 |

\* Data obtained from repurposing tool<sup>79</sup> of cMap database and DrugBank.<sup>31</sup>

#### Experimental validation

Next, we experimentally validated our predictions by targeting genes that have not been reported in the literature to suppress CD4+ T cell proliferation. We selected ten FDA-approved drugs, including at least one target from each disease, based on their association with reactions belonging to pathways we were interested in. Four of the tested drugs indeed resulted in decreased CD4+ T cell proliferation. Genes targeted by these drugs include: COMT (RA, PBC), CPT2 (MS), DPYD (MS), and ACSL3 (PBC). COMT is associated with five different reactions of the tyrosine metabolism (Fig. 7f). The genes CPT2 and ACSL3 both catalyze the production of Long-chain Acyl-CoA in fatty acid biosynthesis from different substrates (Fig. 7c). DPYD catalyzes the production of uracil and thymine from dihydrouracil and dihydrothymine in pyrimidine metabolism. We subjected human CD4+ T cells stimulated with TCR signaling and IL-2 to different doses of drugs for 48 h and 72 h. Their proliferation was assessed by the MTT colorimetric cell proliferation assay (Fig. 8). A reduction of CD4+ T cell proliferation was observed for every chosen drug. Entacapone, targeting the COMT gene, showed significant activity at 48 h and 72 h at 100 μM and 1000 μM (Fig. 8a). EPA, targeting ACSL3, decreased proliferation at the highest dose (1000 μM) at 72 h while the other concentrations did not exhibit a significant effect on proliferation (Fig. 8b). Perhexiline, targeting CPT2, also reduced proliferation at 72 h at 100 μM and 1000 μM (Fig. 8c). Fluorouracil, targeting DPYD, is the only drug that showed an effective impact on CD4+ T cell proliferation at a lower dose (1 μM) at 72 h (Fig. 8d). Interestingly, Entacapone, Perhexiline exhibit a significant enhancement of proliferation at a low dose (1 μM), indicating that these drugs can present a biphasic effect in T cell proliferation. Our experimental validations thus indicate that the perturbation of the activity of our predicted targets can impact CD4+ T cells proliferation in a time and dose-dependent manner.

**Fig. 8:**
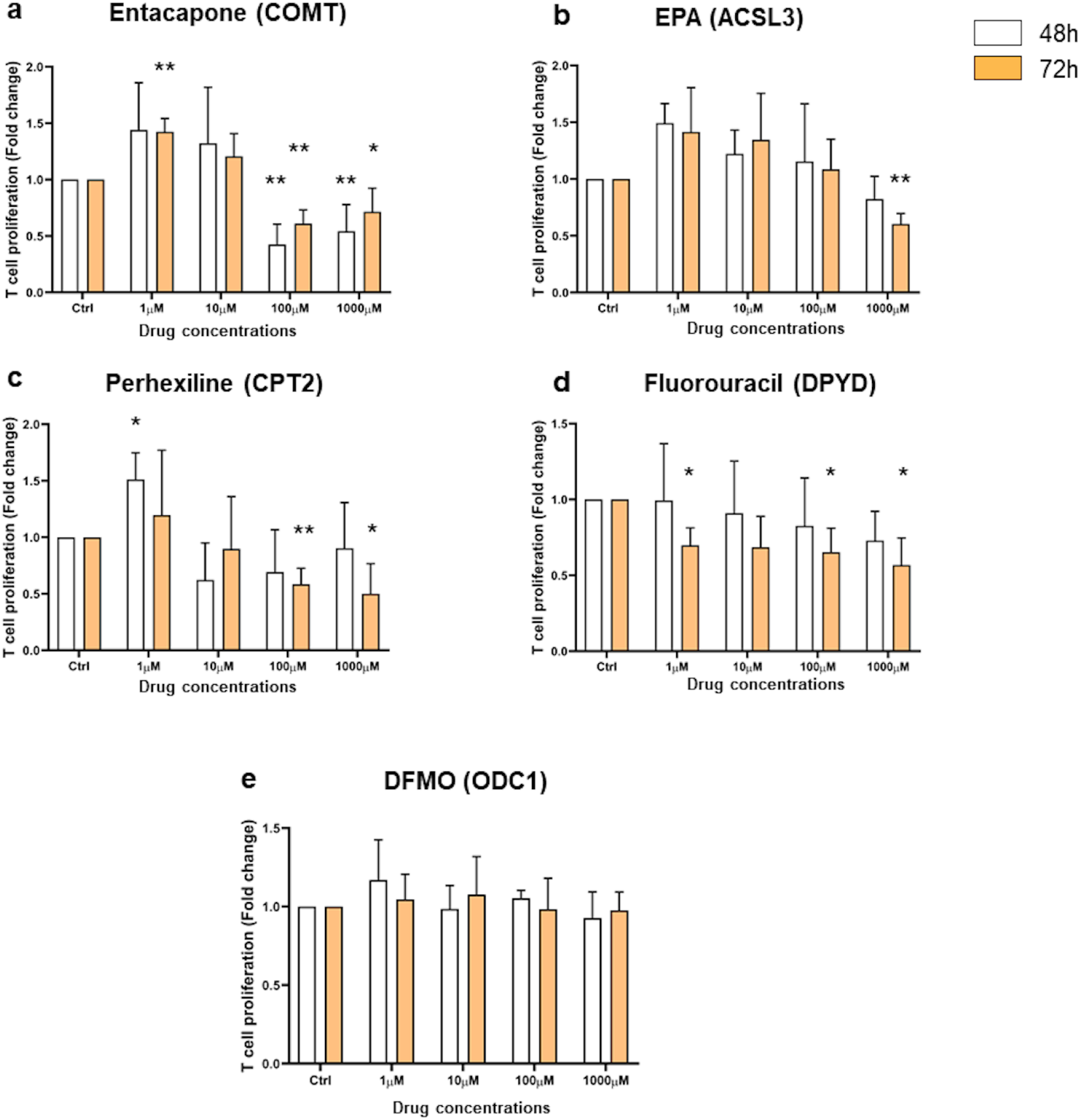
Analysis of CD4+ T cell proliferation response upon drug treatment by MTT assay. CD4+ T cells were exposed to various concentrations of drugs (1 μM, 10 μM, 100 μM and 1000 μM) for 48 h (white bars) and 72 h (orange bars). Drugs’ names (Entacapone, EPA, Perhexiline, Fluorouracil, and DFMO) were indicated on the top of each graph bar with their corresponding targeted gene in parentheses. Cell proliferation is expressed as fold change +/- SEM relative to untreated control cells and is representative of five independent experiments. Statistic significance was only shown for effective concentration and was evaluated using paired t-test, one-tailed (* p < 0.05, **p < 0.005).

Six drugs showed no or opposite effect on the T cell proliferation, including DFMO, Pemetrexed, N6022, Ethyl-pyruvate, Pyrazinamide, and Quercetin. As an example, Fig. 8e shows that DFMO, targeting ODC1 does not impact T cell proliferation with indicating dose and period of time.

Taken together our analysis has identified 68 possible drug targets of relevance to metabolic regulation of autoimmune diseases. We discuss the implications of our results in more detail below.

## Discussion

Our predicted drug targets were classified into two categories: validated, and novel. We considered a target to be validated if previously explored as such in the context of RA, MS and/or PBC. Novel targets, those not previously reported as such for the three diseases we focused on here, were further classified into three subcategories: target genes that are supported by published experimental data, predicted target genes for which no data is currently available, and predicted drug targets for which we provide experimental validation. We will discuss select examples of targets in each of these categories to illustrate ways in which our model and analysis can be used to advance future drug repurposing as well as drug discovery efforts.

Our predictions include three genes (*DHODH*, ACAT1, and *DHFR*) that code for proteins targeted by approved drugs currently used for treatment of autoimmune diseases.^24–28^ A strong example of an already validated drug target predicted by our models is dihydroorotate dehydrogenase (DHODH), a key enzyme in de novo pyrimidine synthesis pathway, and a target of leflunomide, an approved drug for rheumatoid arthritis.^25,29^ DHFR (dihydrofolate reductase) is a well-established oncology drug target, and also as an immunosuppressant and anti-inflammatory target.^30^ Low doses of an FDA-approved DHFR inhibitor methotrexate have been found effective as a treatment for MS, RA, and Crohn’s disease.^27^ Mitochondrial acetyl-CoA acetyltransferase (ACAT1) is a target for FDA approved anti-inflammatory drug sulfasalazine in inflammatory bowel syndrome. Furthermore, this drug is indicated for treatment for rheumatoid arthritis and ulcerative colitis.^31^ Taken together, our models successfully replicated current clinical practice, further strengthening the value of our approach.

Interestingly, a major subcategory of gene targets we identified code for proteins that have not previously been explored for treatment of RA, MS, and PBC. We can now use these insights to formulate novel preclinical and clinical hypotheses. For example, *ABAT*, which encodes the GABA-transaminase enzyme that breaks down γ-aminobutyric acid (GABA; a neurotransmitter), was identified in our analysis as a potential target. While ABAT has not been previously identified as a drug target for RA, we can hypothesize that its inhibition may increase free GABA levels which would, in turn, inhibit CD4+ T cell activation. The relationship between GABA levels and suppressions of CD4+ T cell activation has been previously reported, further suggesting a link between neurotransmission and immune response.^32–34^ There are currently two FDA approved drugs, vigabatrin and phenelzine, that target ABAT. Although we don’t expect that either one of these agents can be repurposed to treat autoimmune disease given that they are an anti-seizure medicine and an antidepressant, respectively, we propose that further analysis of relationships between neurotransmission and immune response offers an interesting targeting opportunity.^24^ Another example is glutathione reductase (GSR), an enzyme that reduces oxidized glutathione disulfide to cellular antioxidant GSH.^35^ It has been shown that inhibition of the de novo GSH synthesis can reduce the pathological progression of experimental autoimmune encephalomyelitis (EAE).^26^ Here, carmustine, a chemotherapy drug, is FDA approved drug that targets GSR, offering a viable starting point for future pre-clinical testing (Table 4). Additional high confidence predictions and target candidates are a group of genes that have been experimentally shown to repress CD4+ T cells upon inhibition. This list includes nicotinamide phosphoribosyltransferase (NAMPT), which we now predict is a drug target for RA. This enzyme is involved in NAD+ synthesis^36^ and was previously explored as a drug target in EAE for MS^37^, melanoma, T cell lymphoma, and leukemia.^38^ Given that two NAMPT inhibitors, GMX1778 and FK-866, are in phase II clinical trials (Table 4), this enzyme represents a target where pre-clinical testing and follow-up may lead to drug repurposing opportunities. Another example worth highlighting is epoxide hydrolase 2 (EPHX2), which converts toxic epoxides to non-toxic dihydrodiols.^35,36^ Its inhibition was reported to result in decreased production of proinflammatory cytokines in preclinical evaluation in inflammatory bowel syndrome.^39^ For EPHX2, an inhibitor GSK2256294A is in phase I clinical trial, indicating that developing drugs for this target may be possible. Moreover, our model implicated enzymes such as pyruvate kinase (PKM), which impacts glycolysis, HMG-CoA reductase (HMGCR), which regulates cholesterol biosynthesis and adenosine deaminase (ADA), which converts harmful deoxyadenosine to not harmful deoxyinosine. Each one of these enzymes has been experimentally linked to T cell proliferation and development,^40–42^ and all three targets have been the subject of previous drug development campaigns (Table 4). ATP Citrate Lyase (ACLY), Catalase (CAT), Farnesyl diphosphate synthase (FDPS), Lanosterol synthase (LSS), Squalene epoxidase (SQLE), and Superoxide dismutase 2 (SOD2) also represent targets we identified. SQLE is involved in cholesterol biosynthesis, and in general agreement with recent reports that inhibiting cholesterol pathways can suppress T cell proliferation.^41^ The loss of SOD2 can increase superoxides and defective T cell development.^43^ For all these targets, either preclinical, clinical or approved inhibitors are available, which we consider encouraging for further study and drug repurposing (see Table 4 for details).

Other predicted genes are part of the TCA cycle (Citrate synthase (CS), Isocitrate dehydrogenase 2 (IDH2)), ribonucleotide biosynthetic processes (Phosphoribosyl pyrophosphate amidotransferase (PPAT), Carbamoyl-phosphate synthetase 2, Aspartate transcarbamylase, and Dihydroorotase (CAD)), and lipid biosynthesis (Fatty acid synthase (FASN)) that are also important for T cell development. As with examples above, many of these potential candidate targets have inhibitors that are in different stages of preclinical and clinical development, and some (like PPAT and FASN inhibitors) have been FDA approved (Table 4). In addition to gene targets with robust or partial existing experimental evidence, we identified 31 novel gene targets for which no evidence currently exists. These genes are involved in glycolysis, TCA cycle, OXPHOS, fatty acid metabolism, pyruvate metabolism, purine, and pyrimidine metabolism, arginine and proline metabolism,and tyrosine metabolism pathways, which are critical for T cell activation and proliferation.^44,45^ We performed *in vitro* experiments to investigate predicted targets without existing published evidence and we tested ten FDA-approved drugs. In four cases of drug targets (COMT, ACSL3, CPT2, and DPYD), the drugs (entacapone, EPA, perhexiline, and fluorouracil, respectively) have inhibited the proliferation of CD4+ T cells in a dose and time-dependent manner. Some drugs displayed a “biphasic effect”, where it encouraged proliferation at a low dose, while inhibiting proliferation at a higher dose. This dose-response effect, called hormesis, is usually an adaptive survival response to toxic agents or stressful environments. Hormetic effects are in general produced by highly conserved evolutionary systems, and characterized by the coordinated activation of molecular signals triggering coherent metabolic responses to maintain cell homeostasis.^46^ Indeed, it is well established that the ability of T cell to modulate their metabolism upon environmental changes is key to preserve their survival and fulfil their functions.^47^ In the case of ODC1, we did not observe any impact on CD4+ T cell proliferation under any tested concentration of DFMO at both 48 and 72 hours. It is difficult to explain this lack of effect without further experimental investigations. It might be that ODC1 is in large excess in our cell cultures, or that the amount of active enzyme is tightly regulated. In addition, each subset of T helper cells is associated with particular metabolic circuits, increasing the difficulty of assessing precise pharmacological strategies to target them. Our computational model is taking into account the precise metabolic requirements for each subset of CD4+ T cells, reducing the space to explore in order to specifically target the essential metabolic pathway in a given cell type.

Collectively, our models and approach led to identification of potential high-value targets for RA, MS and PBC treatment, and proposed several drugs in current clinical use for drug repurposing.

Data integration enabled us to build, refine, and validate high-quality cell type-specific models. While many of the major pathways important for CD4+ T cell activation and proliferation are commonly active across different CD4+ T cell subtypes we tested, the models differ with respect to how these pathways are used for growth. For example, higher activity of the fatty acid oxidation pathway is more important in naïve but not in effector CD4+ T cells that have elevated glycolysis^44^ and fatty acid synthesis pathways. A significant number of essential genes identified from these models overlapped with gene essentiality data obtained from different cell lines.^23^ We are conscious that assessing model performances based on comparison with essential genes defined on different cell lines is not ideal. Properly assessing their accuracy would require CD4+ T cell-specific essentiality data. Integration of disease-associated DEGs with flux profiles under gene knock-out helped us to select disease-specific drug targets. While computational models of signal transduction in CD4+ T cells^48,49^ are available, metabolic models of effector and regulatory CD4+ T cells have not been developed (except for naïve CD4+ T cells^50^). Similar metabolic models were previously used to predict drug targets against pathogens^18^ and complex diseases such as cancers.^51^

While our approach can be generalized for human diseases and used with any-omics dataset, the unavailability of reliable data contributes to some limitations. Because of heterogeneity with respect to time after stimulation with cytokines in the available datasets, constructed models represent inclusive metabolic phenotypes during activation and proliferation for each CD4+ T cell subtype. Thus, time-specific data would be required to study metabolic phenotype at a specific time point in CD4+ T cell development. Furthermore, by integrating gene expression and proteomics data, we have improved the identification of active genes compared with the sole use of gene expression or proteomics datasets. This approach presents certain limitations because of post-transcriptional and post-translational modifications. Integrating more functional data such as enzyme activities and measured metabolic fluxes, could further improve the selection of active reactions within the context of specific models. In addition, the biomass objective function used in our study is not specific to CD4+ T cells. We used a biomass objective function generated for human alveolar macrophages. Differences in growth conditions and the respective sizes of macrophages and CD4+ T cells might impact the ratios of precursors used for biomass objective functions. Since the precursors for biomass production would be the same across different cell types, in the absence of comprehensive data on precursor rations from a specific cell type, the objective function of similar cells is useful for obtaining flux distribution. A specific objective function that considers varying utilization of precursor metabolites (such as glycolysis intermediates) by different CD4+ T cells for biomass production, might further improve the models. However, we have shown that changing objective functions (from the Recon3D to macrophage model) had no significant impact on the constructed models (see Materials and Methods for details). Similarly, reliable disease-specific data were unavailable for specific CD4+ T cell subtypes, therefore, building subtype specific cell metabolic models under disease conditions was not possible. We mitigated this limitation by integrating disease-specific DEGs from sorted CD4+ T cells with models, which resulted in metabolic fluxes relevant to diseases. In the future, with the availability of more disease and cell type-specific data, our integrative approach may further improve these results.

Overall, our integrative systems modeling approach has provided a new perspective for the treatment of RA, MS, and PBC. Moreover, the newly constructed models may serve as tools to explore metabolism of CD4+ T cells. Additionally, our approach is generalizable to other disease areas for which reliable disease-specific data are available, making it a potentially important computational platform for both novel drug target identification and prioritizing targets for drug repurposing efforts.

## Materials and Methods

### High throughput data acquisition and integration

We collected transcriptomics data from the GEO^52^ database and proteomics data.^53^ A total of 121 transcriptomics^54–62^ and 20 proteomics^53^ samples relevant to the CD4+ T cells were selected (*Supplementary Data 1*). Transcriptomics data analysis was performed using the *affy*^63^ and *limma*^64^ R packages. Because we aimed to characterize gene activities instead of gene expression levels, the processed transcriptomics data were discretized (active = 1; inactive = 0) and samples for each cell type were combined. Genes active in more than 50% of the samples in which the probe was present were considered as active (*see Supplementary Methods 2*). Similarly, proteins expressed in more than 50% of samples in the proteomics dataset were considered as active. In the proteomics datasets, protein IDs were mapped to gene IDs.

Next, we integrated activities from transcriptomics and proteomics datasets. First, biological entities that overlapped in both types of data were selected as high-confidence. Second, we found that some genes were expressed in the majority of transcriptomics datasets, but expressed in less than 50% samples of proteomics data. Similarly, some proteins were identified within groups of highly abundant proteins in multiple samples in proteomics datasets but expressed in less than 50% samples of transcriptomics datasets. Such non-overlapping genes were selected as moderate-confidence based on consensus in single types of-omics data (*Supplementary Methods 2.1.1*). Third, moderate-confidence genes exclusively present in the transcriptomics data were added to the overlapping genes if expressed in at least 90% of samples. Fourth, moderate-confidence genes exclusively present in the proteomics dataset were added if their abundance was ranked in the top 25% (fourth quartile) (*Supplementary Methods 2; Supplementary Data 6*). We used these cutoffs to decrease the false negatives while not selecting false positives by removing genes and proteins that are not expressed in any sample either in transcriptomics or proteomics data.

### Cell type-specific genome-scale metabolic model reconstruction

We used the GIMME^65^ method (in COBRA toolbox) to construct the metabolic models of different CD4+ T cells (naïve, Th1, Th2, Th17). The inputs for GIMME were the generic human Recon3D^66^ (as a template) and gene expressions based on integrated multi-omics data. The template Recon3D was modified prior to constructing CD4+ T cell-specific metabolic models. These modifications included gene-protein-reaction (GPR) associations (all genes associated with a reaction written using AND and OR operators), media conditions, and reaction directionality. In the original Recon3D, GPRs used transcript IDs. Because our data included gene IDs, we mapped the transcript IDs to the gene IDs. For naïve and effector cells, different types of media condition were selected based on nutrient preference information obtained from the literature (*Supplementary Methods 2.1.2*). In addition, new reactions involved in the biomass objective function were added, and some reactions were removed as described below (See also *Supplementary Methods 2*). The transcriptomics and proteomics data have information about genes/proteins instead of transcript variants. To map the data obtained for genes, we updated transcript IDs provided in Recon3D to Entrez gene IDs. A total of 1892 genes were included in the modified Recon3D model. Furthermore, because different CD4+ T cells have different nutrient uptake preferences, we used two types of media conditions (one for each naïve and one for all effector T cells), shown in *Supplementary Methods 2.1.2.4*. For all cell subtypes, in addition to the basal metabolites (freely available, i.e. H2O, O2, H, O2S, CO2, Pi, H2O2, HCO3, H2CO3, and CO), glucose, glutamine, and other amino acids were set as open (but tightly constrained) for uptake. The major difference in media condition was the presence of fatty acids in the naïve model. Furthermore, during the refinement of the CD4+ T cell models, the directionality of some reactions was updated based on the Recon 2.2.05 model^67^ and the MetaCyc database (*Supplementary Methods 2.1.2.5* and *2.2.3*). Because of the lack of CD4+ T cell-specific data, the biomass objective function was adopted from the macrophage model iAB-AMØ-1410^68^ and added to the Recon3D.

For each subtype, we constructed three models based on transcriptomics, proteomics, and integrated (transcriptomics and proteomics data) datasets. A comparison of these models is provided in *Supplementary Figure 15* and details can be found in *Supplementary Methods 2*. The models constructed with integrated data were selected for further analysis. Additionally, to investigate the effect of biomass objective function on constructed models, we built two models using biomass objective functions from (1) Recon3D and (2) iAB-AMØ-1410 models. The reactions in output models generated based on each biomass function were compared. The models based on the two objective functions were not significantly different (*Supplementary Figure 16*) with respect to the numbers of reactions. Biomass objective function from iAB-AMØ-1410 consists of a few extra precursors that predicted better fluxes through fatty acid pathways. The literature has shown that effector CD4+ T cells synthesize fatty acids, whereas naive CD4+ T cells exhibit fatty acid oxidation. We compared the fluxes of models created with objective functions from Recon3D and iAB-AMØ-1410. The models created with biomass objective function adopted from iAB-AMØ-1410 had more reactions carrying non-zero flux in the fatty acid pathways (than the models created with Recon3D biomass objective function). Therefore, models that are constructed based on biomass reaction adopted from iAB-AMØ-1410 were used in subsequent analyses. Models were further reduced by removing the dead-end reactions. Reactions in the models are distributed across different compartments including extracellular, cytoplasm, mitochondria, nucleus, Golgi apparatus, lysosome, and endoplasmic reticulum. The models were investigated to perform basic properties using leak test, gene deletion and further refined in an iterative manner. To examine leaks, we simulated the models with all the exchange reactions closed and analyzed all the reactions individually for non-zero flux. If the models were producing metabolites, the mass imbalance was checked and fixed. We used gene deletion analysis to check if the model was able to predict gene essentiality. Because effector T cells are highly glycolytic, deleting glucose transporters should result in reduced growth. We used this as a reference to check that the model was behaving correctly. Furthermore, we investigated if inhibiting acetyl-CoA carboxylase (ACC1) — which was experimentally observed as essential for CD4+ T cell function — resulted in altered growth. These analyses helped us identify problematic reactions that were corrected based on Recon2.2.05 — a manually curated model for mass charge balancing and reaction directionality — and the MetaCyc database. Refined models were then subjected to 460 metabolic tasks that were used with the Recon3D model and included in *Test4HumanFctExt* function in COBRA (*Supplementary Data 7*). The constructed models were simulated using Flux Balance Analysis (FBA) and Flux Variability Analysis (FVA). The final numbers of metabolites and reactions are presented in Table 1. These models were named as TNM1055 (naïve model), T1M1133 (Th1 model), T2M1127 (Th2 model), and T17M1250 (Th17 model). The models encoded in SBML can be found as *Supplementary Dataset 1*. They have also been submitted to BioModels database^69^ under accessions MODEL1909260003, MODEL1909260004, MODEL1909260005, MODEL1909260006.

### Model validation

Models were validated based on literature knowledge related to active pathways in proliferating and differentiated CD4+ T cells. CD4+ T cell-specific metabolic functions were searched in the literature using PubMed.^70^ Naïve CD4+ T cells tend to have low energy demands, and mainly rely on *fatty acid β-oxidation*, oxidation of pyruvate and glutamine via the *TCA cycle*.^71^ On the other hand, the high bioenergetics demand in effector cells is met by shifting *OXPHOS* to *glycolysis* and *fatty acid oxidation* to *fatty acid synthesis*.^45^ Furthermore, similar to cancer cells, proliferating effector CD4+ T cells convert lactate from pyruvate by lactate dehydrogenase enzyme.^45^ Thus, we obtained the flux distribution of metabolic pathways under wild type conditions using Flux Balance Analysis (FBA) and searched the non-zero fluxes through the aforementioned pathways in all the models. Flux maps were created using Escher web application (https://escher.github.io/#/).^72,73^ It has also been observed previously that deficiency in glucose and glutamine impairs CD4+ T cell activation and proliferation.^74,75^ We performed this experiment *in silico*, whereby we varied the flux through exchange reactions of glucose (EX_glc[e]) and glutamine (EX_gln_L[e]) in the models and analyzed the effect on growth rate.

### Comparison of essential genes predicted by the models and identified in different cell lines

To predict gene essentiality, we knocked out model genes to predict their effect on the growth rate. This was performed using *singleGeneDeletion* in the COBRA toolbox using the Minimization of Metabolic Adjustment (MoMA) method.^76^ Because of the unavailability of CD4+ T cell-specific data, predicted essential genes were compared with experimentally identified essential genes in humans from different cell lines. The data for experimentally tested essential and nonessential genes for human were obtained from the OGEE database.^23^ In this database, the essentiality data for humans was compiled using 18 experiments across various cell lines that include RNAi based inhibition, CRISPR, and CRISPR-CAS9 systems. To investigate how many model-predicted essential genes are also essential in other cell lines, predicted essential genes were compared with experimentally observed essential and conditionally essential genes reported in the OGEE database. Essential and conditionally essential genes were merged together. Additional validations of models using CD4+ T cell-specific essential genes can be found in *Supplementary Methods 3*.

### Mapping drug targets

The developed models were used to predict potential drug targets for autoimmune diseases in which effector subtypes have been found hyperactive.^77,78^ Therefore, a reasonable drug target should downregulate effector CD4+ T cells. Among the metabolic genes of selected models, we first identified targets of existing drugs. The drugs and their annotations including target genes were imported from The Drug Repurposing Hub^79^ in the ConnectivityMap (CMap) database.^80^ All withdrawn drugs and their annotations were first removed. In this list, the gene symbols of target genes of drugs were converted to Entrez IDs. Next, we searched Entrez IDs from CMap data in the genes of metabolic models. For each mapped gene in the model, the drugs were listed.

### Metabolic genes differentially expressed in autoimmune diseases

The lack of reliable data from specific CD4+ T cell subtypes involved in autoimmune disease conditions led us to utilize patients’ data (case-control studies) available for autoimmune diseases that were collected from peripheral CD4+ T cells. Datasets GSE56649^81^ (rheumatoid arthritis), GSE43591^82^ (multiple sclerosis), and GSE93170^83^ (primary biliary cholangitis) were obtained from the GEO database. Raw data files were processed using the *affy* and *limma* packages^63,64^ in Bioconductor/R. The *limma* package was used to identify DEGs between patients and healthy controls. For significant differential expression, selective cutoffs of fold-changes were used with adjusted P-values < 0.05. For differentially expressed genes, we used a two-fold cutoff. A cutoff of 1.5-fold was used for datasets where two-fold resulted in a very low number to zero differentially expressed metabolic genes.

### Perturbation of metabolism and perturbation effect score (PES)

In metabolic models, the knockout of genes that are targets of existing drugs was performed in the COBRA toolbox using MoMA.^76^ For each knockout, we investigated the change in fluxes regulated by DEGs in diseases. The change in fluxes was computed using flux ratios of perturbed flux/WT flux, and all fluxes that are affected by each perturbation were calculated. We counted fluxes regulated by upregulated genes that are decreased or increased after perturbation (UpDec and UpInc) as well as fluxes regulated by downregulated genes that are decreased or increased after perturbation (DownDec and DownInc). The total number of fluxes for each perturbation also include upregulated genes that were unchanged after perturbation (UpUnc) as well as downregulated genes that were unchanged after perturbation (DownUnc) (see also *Supplementary Methods 7*). For each perturbed gene, a perturbation effect score (PES) was calculated as:

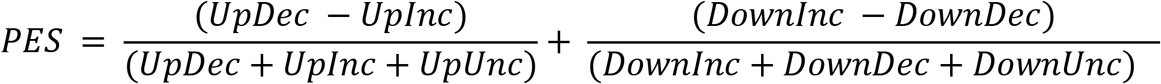

Next, for each disease and model combination, the ranks of PES were computed. The gene with the highest PES obtained the top rank and the one with the minimum PES obtained the lowest rank. For each disease, we prioritized drug targets by utilizing their ranks across all models. The PES ranks in each model were first transformed into Z-score as:

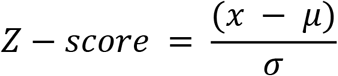

where *x* is a PES rank, *μ* is the mean of PES ranks in a model for one disease, σ is the standard deviation of PES ranks obtained by a model for one disease. For each disease type and each gene, Z-scores across four models were summed up to calculate an aggregated Z-score. Genes were ranked based on minimum to maximum aggregate Z-scores.

### Pathway enrichment analysis

For biological processes enrichment analysis, we used DAVID V6.8,^84^ and STRING database^85^ together with Gene Ontology biological processes,^86^ KEGG pathways,^87^ and Reactome pathways.^88^ A cutoff of 5%^89,90^ False Discovery Rate (FDR) and P-value < 0.05 were used for significant enrichment. The pathway maps used in Fig. 7 were generated with yED graph editor software.

### Experimental Validation

#### Cells culture

We used frozen vials of peripheral blood mononuclear (IXCells Biotechnologies) from healthy donors. CD4+ were purified by negative selection using magnetic human CD4+ T cells nanobeads (MojoSort, Biolegend) according to the manufacturer’s protocol. For cell activation, anti-CD3 (clone OKT3, Biolegend) was coated on plates at 4 μg/ml overnight in PBS at 4 °C. Cells were then cultured in X-Vivo media (Lonza) supplemented with 2 μg/ml of anti-CD28 (clone CD28.2, Biolegend) and with 10 ng/ml of recombinant IL-2 (Peprotech) for 7 days. Half of the media was renewed every 2-3 days by adding fresh media supplemented with 2 μg/ml of anti-CD28 and 10 ng/ml IL-2.

#### Drug treatment

We purchased alpha-difluoromethylornithine (DFMO) and eicosapentaenoic Acid (EPA) from Cayman Chemical. DFMO was resuspended in water at 50mM while EPA was already resuspended in ethanol at 826 mM. Perhexiline, Entacapone, and Fluorouracil were obtained from Tocris and dissolved in DMSO to make stock solution at 12 mM, 200 mM, 198 mM, respectively. For drug treatment, CD4+ T cells were seeded at 50000 cells in 96 round bottom wells in culture media supplemented with anti-CD3, anti-CD28 and IL-2. Before incubation with CD4+ T cells, drugs were diluted in culture media with concentration ranging from 1 μM to 1000 μM and incubated for 48 h or 72 h.

#### MTT assay

We assessed T cell proliferation using the TACS MTT proliferation assay (R&D systems). Briefly, a tetrazolium salt solution (10 μl) was added to each well, and the plate was incubated at 37 °C for 4 h. After incubation, 100 μl of stop solution was added to each well and incubated overnight before absorbance measurement. Cell proliferation was read at 48 h and 72 h post-drug treatment. The absorbance was measured at 570 nm using the BioTek microplate reader instrument (BioSPX). We corrected cell absorbance readings using cells treated with DMSO or the media controls.

#### Statistical analysis

The CD4+ T cell proliferation upon drug treatment was analyzed statistically with a paired t-test, one-tailed for five independent experiments. All data are presented as mean plus or minus standard error of the mean (SEM) and analyzed using GraphPad Prism software. The fold change of cell proliferation-cultured was calculated using untreated cells as 1.

## Competing Interests

The authors have declared that no competing interests exist.

## Acknowledgements

This work was supported by NIH grant 1R35GM119770-04 to T.H. and by the Systems Biology Grant of the University of Surrey to M.B.

## Author Contributions

T.H. and B.L.P. conceived the study. B.L.P., M.B., and T.H. designed the study. B.L.P., M.B., and T.H. defined the pipeline to integrate-omics data, and B.L.P implemented the pipeline. B.L.P. and B.L. performed literature mining. B.L.P., B.L., R.M., A.C., and S.T. collected the data. B.L.P., B.L., R.M., and A.C. constructed the models. A.R.S. performed refinement of the constructed models. B.L.P., B.L., R.M., A.C., S.T., M.B., and T.H. analyzed the data. R.A. performed the experimental work and analyzed the experimental results. B.L.P., M.B., and T.H. wrote the manuscript. M.B. and T.H. supervised the study.

## Availability of data and materials

The models generated in this study are available as SBML files in the *Supplementary Dataset 1* and can also be accessed from the biomodels database under accession MODEL1909260003, MODEL1909260004, MODEL1909260005, MODEL1909260006. The scripts used in this pipeline are available from the corresponding author on reasonable request.

## Notes

### Competing Interest Statement

The authors have declared no competing interest.

